# Role of CRF receptor 1 antagonism in the bed nucleus of the stria terminalis of male rats after intermittent social defeat stress

**DOI:** 10.1101/784488

**Authors:** Mailton Vasconcelos, Dirson J. Stein, Matheus Gallas-Lopes, Luane Landau, Luiza Behrens, Lucas Albrechet-Souza, Klaus A. Miczek, Rosa Maria M. de Almeida

**Affiliations:** Instituto de Psicologia, Universidade Federal do Rio Grande do Sul, Porto Alegre, RS – Brazil; Department of Physiology, Louisiana State University Health Sciences Center, New Orleans, LA – USA; Departments of Psychology and of Neuroscience, Tufts University, Boston, MA – USA

**Keywords:** social stress, social approach, anxiety, corticotropin-releasing factor receptor 1, bed nucleus of the stria terminalis

## Abstract

We recently demonstrated that the experience of brief episodes of social defeat caused impairments in social behaviors. Moreover, we provided evidence that the antagonism of corticotropin-releasing factor binding protein (CRFBP) in the bed nucleus of the stria terminalis (BNST) restored social approach in stressed animals. This study aimed to test the relation between corticotropin-releasing factor receptor type 1 (CRFR1) located in the BNST and the establishment of social stress-disrupted behaviors in rats submitted to social defeat in the resident-intruder paradigm. Animals were tested for sweet solution preference, subjected to the elevated-plus maze (EPM), and to the social interaction three-chamber test. Social behavior was tested after BNST drug infusions. The drug used in this study was a CRF receptor 1 antagonist, CP376395 (CP), administered in two doses: 50 ng/0.20 μL/side, and 500 ng/0.20 μL/side. Saline solution was used as vehicle and administered 0.20 μL/side. Socially stressed animals (n = 11) did not differ compared to control animals (n = 11) in the EPM. Stressed animals displayed impaired social behavior, represented by a decrease in time spent in the interaction zone. The lower dose (CP 50 ng/0.20 μL/side) administered intra-BNST restored social behaviors in stressed animals. On the other hand, the higher dose of the CRFR1 antagonist (CP 500 ng/0.20 μL/side) induced social avoidance in rats without a history of agonistic confrontations. These findings implicate BNST CRFR1 signaling in the modulation of social behaviors in rats given the choice to explore an unfamiliar conspecific.

## 1. Introduction

The BNST is an important extra-hypothalamic site of action for the corticotropin-releasing factor (CRF) in potentiating stress responses, mainly those that are linked to anxiety-like behaviors [1]. Pharmacological manipulations of the CRF activity in the BNST of rodents demonstrated its relevance in startle responses [2,3], anxiety-like behavior, place aversion [4], and reinstatement of drug seeking behaviors [5]. Many of these findings support the hypothesis that blockade of CRF actions will alleviate symptoms of depression, anxiety, and drug addiction. Since the first characterization of CRF, the identification of CRF antagonists pharmacological properties was a major goal [6]. CRF and its family of ligands transduce neural and endocrine signals by binding to two major CRF receptor types, CRFR1 and CRFR2 [7]. Additionally, the actions of CRF and related peptides can be modulated by the CRFBP [7].

Recently, we demonstrated that social approach was restored after local infusions of the CRFBP antagonist, CRF fragment 6–33, into the BNST of socially stressed animals [8]. These experiments demonstrated the involvement of CRF signaling in the development of stress-induced effects after the experience of brief episodes of social defeat stress in male rats. Nevertheless, it remains unclear which CRF receptor subtype is involved in this stress-induced disruption of social behavior. The current experiment was conducted to evaluate the behavioral effects of CRFR1 antagonist microinjection into the BNST of rats submitted to intermittent social defeat stress.

## 2. Materials and Methods

### 2.1 Animals

Adult male Wistar rats, termed “intruders”, were obtained at approximately postnatal day (PND) 50 and housed for 2 weeks of habituation at the Animal Experimental Unit of Hospital de Clínicas de Porto Alegre, RS, Brazil before starting the experiments. At PND 70, rats were singly housed in custom-built acrylic cages (15 × 25 × 20 cm). Separate male Wistar rats, weighing 577.59 ± 41.12 g, with a reliable history of aggressive behavior, were housed in pairs with a sterile female Wistar rat in large custom-built acrylic cages (46 × 71 × 46 cm). These aggressive rats were designated as “residents”. Both cohorts of animals were kept in separate rooms with controlled environmental conditions: 21 ± 1° C of temperature, 40-60% of humidity and controlled 12 h light-dark cycle (lights on at 7 am). The cages were lined with sawdust bedding and animals had free access to food and water. A total of *N* = 57 rats was used in this study (11 aggressive males, 11 females paired with the aggressive males, 5 ovariectomized females for the social approach test, and 30 male intruders). This study was supervised and approved by the *Comitê de Ética para Uso de Animais do Hospital de Clínicas de Porto Alegre*, and executed following the Guide for the Care and Use of Laboratory Animals [9] and Brazilian law 11.794/2008.

### 2.2 Drug

The drug used in this study was a CRFR1 specific antagonist, CP376395 (CP) obtained from Bio-Techne (Minneapolis, MN, USA). CP was diluted in 0.9% sterile saline solution at final concentrations of 250 and 2500 ng / μL.

### 2.3 Experimental design

The stress protocol was conducted following procedures established by Miczek and colleagues [10–12]. Intruder rats were subjected to four intermittent episodes of social defeat stress over the course of 10 days or served as contemporary unstressed controls. After concluding the stress protocol, rats were tested for sweet solution preference (PND 85), went through stereotaxic surgery (PND 90-91), a second sweet solution preference test (PND 95), followed by anxiety-like and risk assessment behaviors in the EPM test (PND 96). Finally, the animals received bilateral intra-BNST microinjection of either saline solution or CP in two different doses (50 ng/0.20 μL/side and 500 ng/0.20 μL/side) and were tested in the three-chamber social interaction test (PND 97, 99, and101).

### 2.4 Intermittent social defeat stress

Intruders went through a regimen of intermittent social defeat stress of four sessions with 72h interval between them [8,12]. At the beginning of this resident-intruder protocol, the females were removed from the large resident cages, and intruders were exposed to an unfamiliar resident in a protected home cage. This initial procedure lasted 10 min in order to instigate territorial behavior and heighten aggressiveness of the residents. The intruder was removed from its protective cage and was directly confronted by the aggressor. There were three criteria to end this fighting session: after the intruder displays a supine posture for 5s, or 5min after the resident’s first biting attack, or after 5 bites, whichever occurs first. After the confrontation, the intruder was placed back into the protective cage within the resident’s home cage for additional 10 min [8,12]. Control rats were handled and weighed, without being submitted to the stress protocol.

### 2.5 Sweet solution preference

The sweet solution preference was used to test animals for reduction in sensitivity for positive reinforcement. The reduction in social approach was chosen as the main outcome of the study. To test whether this stress-induced effect would be due to a depressive-like state we monitored the preference for sweet solution of animals in PND 85 and 95. To perform the test, rats were removed from their home cages, transferred to a behavioral testing room, and assessed for preference to a sweetened drink (0.8% saccharin and sodium cyclamate; Zero·Cal, Hypermarcas; São Paulo, SP, Brazil) or water during one hour in a test cage (24 × 38 × 15cm). The results of these tests were compared with the animals’ previous sweet solution preference before exposure to the stress protocol. To verify the baseline preference, at PND 71, rats were exposed overnight to one bottle containing the sweet solution and another bottle containing tap water. In the four consecutive days, PND 72–75, rats were given a daily 2-bottle choice (sweet solution and water) for one hour. The position of the bottles was switched between trials to counteract side preference. Animals were not food or water deprived, before or after trials. It was used an empty “drip” cage to serve as control for evaporation and spillage due to handling of bottles. The weight of the bottles was used to measure fluid consumption. The preference was calculated by dividing the total amount of sweet solution consumed by the total fluid intake (water plus sweet solution) [8,13].

### 2.6 Stereotaxic surgery

Stereotaxic surgery was conducted between PND 90-91, at least five days before drug infusion. Rats were anesthetized with isoflurane. Tramadol (20 mg/kg, i.p.) and bupivacaine (8 mg/kg, intradermal) were used as pre-surgical analgesia and local anesthesia respectively. The intruders were mounted on a stereotaxic frame (Kopf Instruments; Tujunga, CA, USA) and guide cannulae were bilaterally implanted into the BNST using the coordinates from bregma: AP = 1.0; ML = +/−2.5; DV = −8.0, with a 10° angle [8,14]. Stainless steel screws and dental cement were used to fix guide cannulae, additionally custom made dummy cannulae were used to seal the guides. Post-operative analgesia was conducted with dipyrone (500 mg/kg, i.p.), and subsequent pain control was provided with tramadol (20 mg/kg, i.p.), every 12h for 2 days.

### 2.7 Elevated-plus maze test

At PND 96, rats were submitted to the EPM test. The EPM apparatus comprised two open arms (50 × 10 cm) and two closed arms (50 × 10 × 50 cm) connected to a central platform (10 × 10 cm). Arms of the same configuration were opposite to each other, and the entire apparatus was elevated 50 cm from the floor [15]. The animals’ behaviors were video recorded from above. Rats were individually placed in the center of the apparatus facing a closed arm and allowed to freely explore the entire plus-maze for 5 min. The behaviors were automatically recorded using a behavioral tracking system (ANY-Maze, Wood Dale, IL, USA). The software provided measurements of the number of entries and time spent in each part of the maze. An arm entry was defined by the presence of all four paws of the animal in the specific arm of the maze. Additionally, risk assessment behaviors (head-dipping, stretch-attend postures, and rearing) were coded manually using BORIS event-logging software [16].

### 2.6 Microinjection

Intra-BNST microinjections occurred at PND 97, 99, and 101 starting 5 minutes before social approach test. Drug administrations were counterbalanced in a way that saline solution and CP doses were tested on the same day in different animals. Each animal received one bilateral microinjection per day, but they were tested in all drug conditions throughout the three days of social interaction. All procedures occurred during the first four hours of the dark phase.

### 2.7 Social approach test

Animals were tested for social approach in the three-chamber social test at PND 97, 99, and 101. The apparatus consisted of an acrylic box (40 × 50 × 120 cm) with walls separating the box into three chambers [8]. There were central openings that, once the pair of removable walls were lifted, allowed the animal to move freely throughout the apparatus. The test started with the intruder confined to the middle chamber for 5min. Two acrylic cages, identical to the intruders’ home cage (15 × 25 × 20cm), were each one placed in each of the lateral chambers to perform the test. An unfamiliar ovariectomized female was placed inside of one of the cages in one of the side chambers. One cage was left empty in the opposite side chamber. The central walls were opened, and the intruders could explore all three chambers over a period of 10 min. The small cages provided similar protection, as in the social defeat protocol, from any aggressive interaction between the animals. During trials, the female, the location of the female, and the empty cage were alternated between the upper and lower corners of the side chamber in consecutive trials. Sessions were video recorded and behaviors were coded using event-logging software [16]. Frequency and duration of entries into the side chambers containing the unfamiliar female (interaction zone) or the empty cage (object zone) were measured for 10 min [17]. The apparatus was cleaned with 70% alcohol between trials. Animals were tested after microinjection, with one session a day, with a 48h-interval between the each of the three sessions.

### 2.8 Histological confirmation

On the day after the end of behavioral tests, rats were deeply anesthetized with an overdose of isoflurane (5% for more than 3 min) and perfused with 0.9% saline followed by 4% formaldehyde solution prior to brain removal. The fixed brains were sliced into 40 μm coronal sections using a cryostat (Microm HM 525, Thermo Fisher Scientific, Waltham, MA, USA). Slices were mounted on glass slides and stained with 0.01% Toluidine Blue solution. Injector placements were verified using light microscopy, according to a rat brain atlas [14]. Diagrammatic representations of bilateral BNST injection sites are shown in Figure 2a.

**Figure 1.**
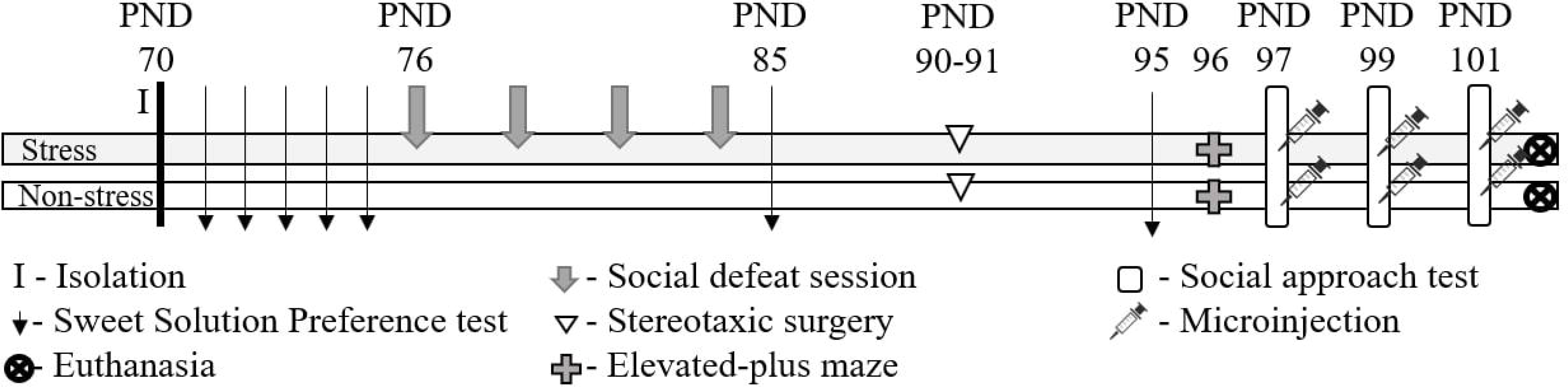
Experimental procedure. Stress: rats exposed to intermittent episodes of social defeat; Non-stress: control animals; *N* = 11 rats per group.

**Figure 2.**
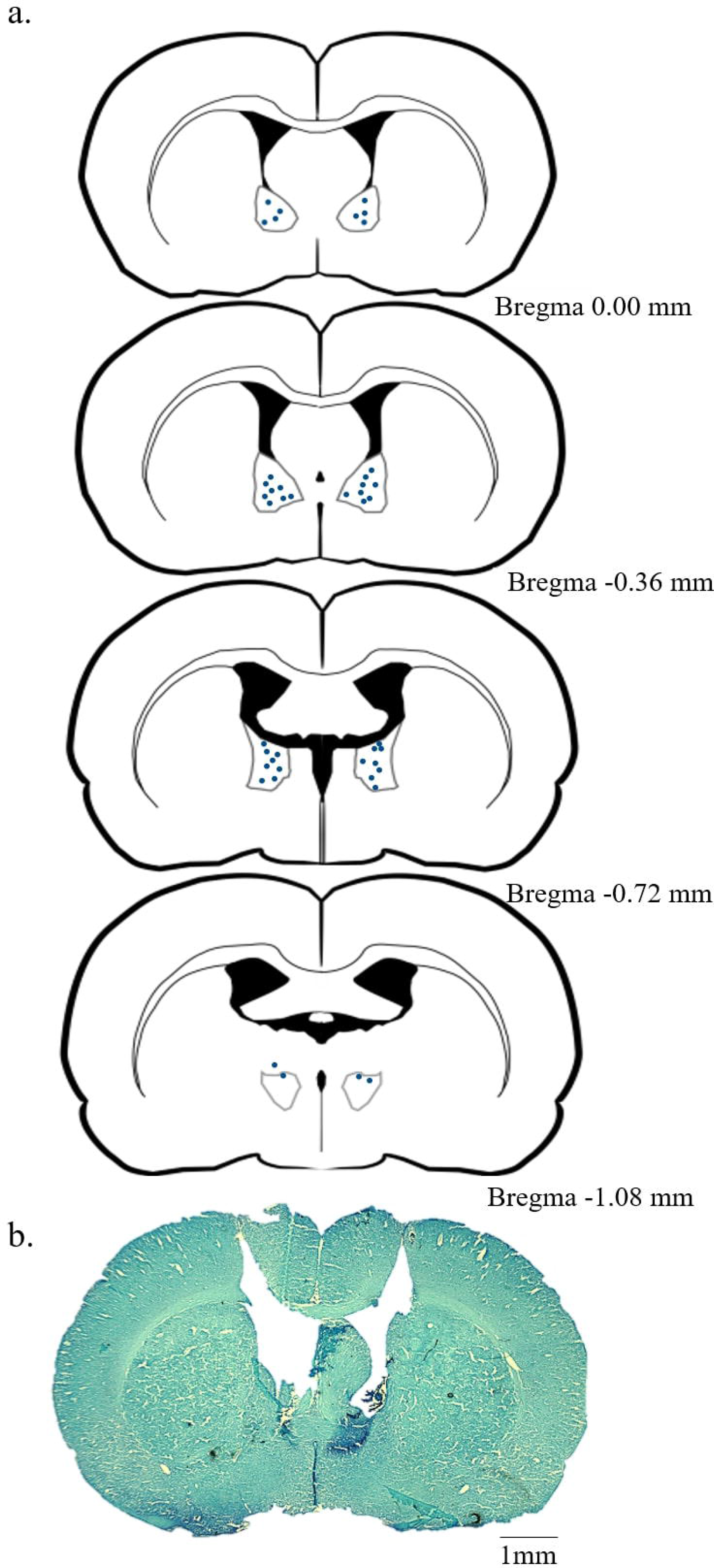
Diagrams and photomicrograph of histological confirmation procedures a. Correct cannulae placements into the bed nucleus of the stria terminalis; b. Representative photomicrograph after toluidine blue staining.

### 2.7 Statistical analysis

To assess the effects of social stress on weight gain and preference for sweet solution, a two-way repeated measures ANOVA was conducted (GraphPad Prism 6 for Windows, GraphPad Software Inc., La Jolla, CA, USA). Stress and non-stress were set as conditions, and baseline and two additional time point measurements were set as sessions. Independent-samples *t*-tests were conducted to compare frequency and time spent in open and closed arms of the EPM, as well as the frequency of risk-assessment behaviors, differentiating non-stress and stress as conditions. To assess social approach, the frequency of entries and time spent in either interaction zone (IZE and IZT), center (CeE and CeT) and object zone (OZE and OZT) were analyzed. We also assessed the latency to first enter the interaction zone and computed the average time spent in the interaction zone per frequency of visit. A two-way repeated measures ANOVA was performed to analyze the effect of CP microinjection and social defeat on social approach. Stress and non-stress were set as conditions and saline and two different doses of CP microinjections (50 ng/0.20 μL/side and 500 ng/0.20 μL/side) were set as treatments. When resulting in a significant overall effect, *post hoc* multiple comparisons were performed using Bonferroni test. The statistical significance was set at *p* < 0.05. t-tests were conducted to compare stress vs. non-stress groups for the frequency and time spent in the interaction zone under saline microinjection. Data were expressed as mean ± SEM.

## 3. Results

Rats with injector tracks that did not reach the BNST, and rats that did not complete three days of drug treatment for reasons related to health conditions were excluded from the analysis (n = 8; 14.03% of the total sample size). In cases where the behavior of the animal was recorded but the data could not be extracted, an imputation technique was implemented to replace any missing values with the mean of the variable computed from all other cases (24 values, 2.66% of all data).

The sweet solution preference test was conducted to investigate the effect of stress on hedonic behaviors. The protocol is relatively non-stressful and non-invasive, allowing multiple testing without further influence on the animal’s behavior. The present results demonstrated no differences in preference for a sweet solution after the acute experience of social defeat and no differences 10 days after the last day of stress exposure. There were no significant effects on preference for sweet solution across sessions as revealed by two-way repeated measures ANOVA (*F*(2, 40) = 0.88). Rats did not show significant differences in preference between conditions (*F*(1, 20) = 0.01), nor across time (*F*(2, 40) = 1.50) (Table 1).

**Table 1.**
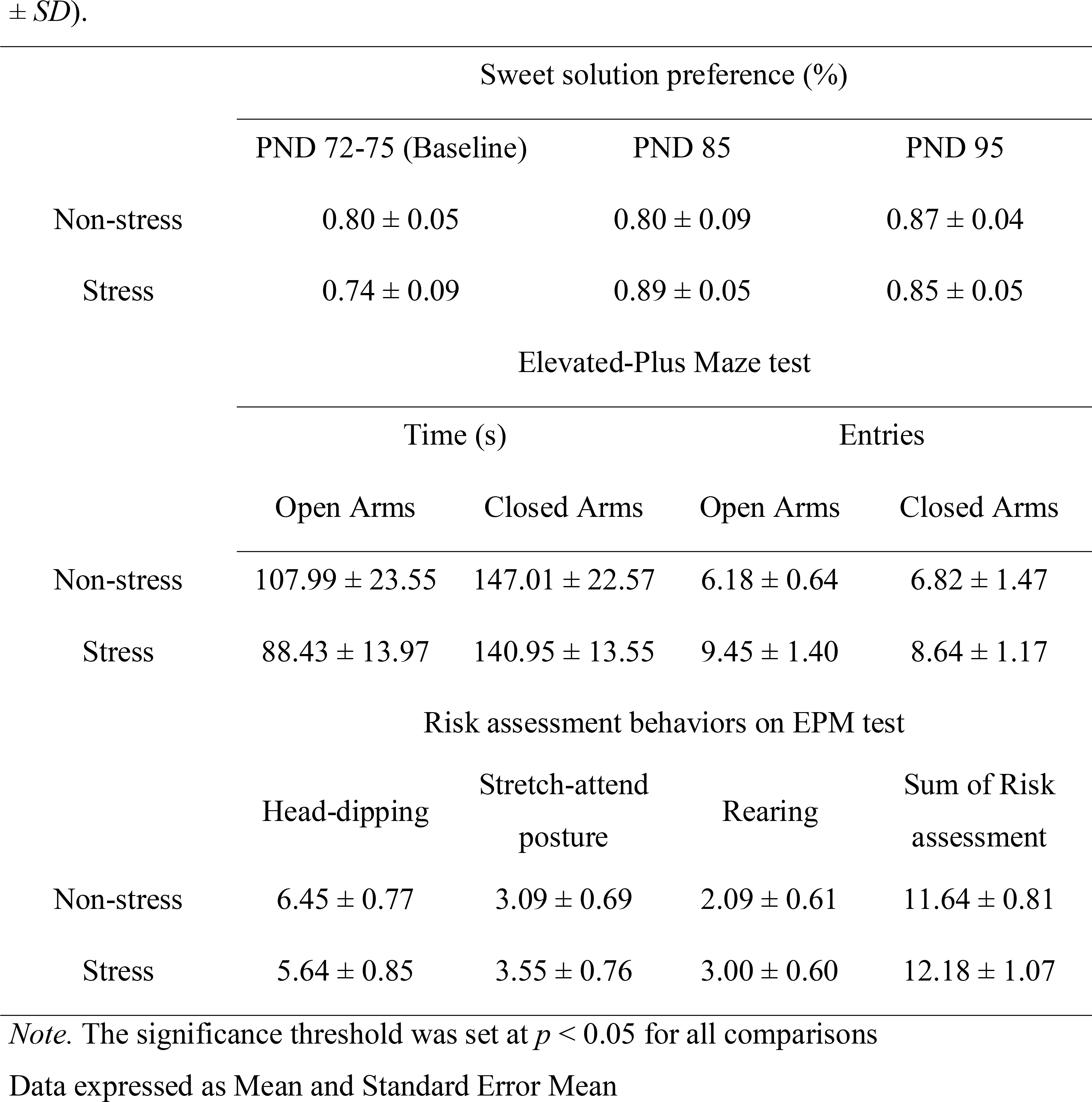
Data of preference for a sweet solution (*M* ± *SEM*) during baseline and two additional measurements; data of behaviors displayed in the Elevated-Plus Maze test (*M* ± *SD*).

|  | Sweet solution preference (%) |  |  |  |
| --- | --- | --- | --- | --- |
|  | PND 72-75 (Baseline) |  | PND 85 | PND 95 |
| Non-stress | 0.80 ± 0.05 |  | 0.80 ± 0.09 | 0.87 ± 0.04 |
| Stress | 0.74 ± 0.09 |  | 0.89 ± 0.05 | 0.85 ± 0.05 |
|  | Elevated-Plus Maze test |  |  |  |
|  | Time (s) |  | Entries |  |
|  | Open Arms | Closed Arms | Open Arms | Closed Arms |
| Non-stress | 107.99 ± 23.55 | 147.01 ± 22.57 | 6.18 ± 0.64 | 6.82 ± 1.47 |
| Stress | 88.43 ± 13.97 | 140.95 ± 13.55 | 9.45 ± 1.40 | 8.64 ± 1.17 |
|  | Risk assessment behaviors on EPM test |  |  |  |
|  | Head-dipping | Stretch-attend posture | Rearing | Sum of Risk assessment |
| Non-stress | 6.45 ± 0.77 | 3.09 ± 0.69 | 2.09 ± 0.61 | 11.64 ± 0.81 |
| Stress | 5.64 ± 0.85 | 3.55 ± 0.76 | 3.00 ± 0.60 | 12.18 ± 1.07 |

*Note.* The significance threshold was set at $p < 0.05$ for all comparisons
Data expressed as Mean and Standard Error Mean

Deficits in social interaction have been classically interpreted as a sign of anxiety-like social behavior [18]. We decided to test whether exposure to an aggressive resident would generate anxiety-like behavior measurable in the EPM test. Independent-samples *t*-tests were conducted to compare frequency and time spent in open and closed arms of the EPM. There was no significant difference in the frequency of entries and time spent into the arms of the maze between the conditions (open arm time: *t(*20) = 0.71; and closed arm time: *t(*20) = 0.23; open arm frequency: *t(*20) = 1.50; closed arm frequency: *t(*20) = 0.97). There was no significant difference in the frequency of any of the risk-assessment behaviors between conditions (head dipping: *t(*20) = 0.50; stretch-attend posture: *t(*20) = 0.31; rearing: *t(*20) = 0.75; and sum of all risk-assessment behaviors: *t(*20) = 0.29). These results suggest that exposure to intermittent social defeat stress produced no effects on behaviors related to anxiety displayed in the EPM apparatus (Table 1).

Based on our previous findings [8], this study relies on the premise that stressed animals present reduced interest in approaching a conspecific in the three-chamber social test. This was tested by comparing behaviors displayed in the interaction zone between conditions, stress/non-stress, under intra-BNST saline microinjection. Rats from the non-stress condition spent significantly more time in the interaction zone (IZT: 249.90, SE± 13.20) compared to rats in the stress condition (IZT: 199.70 ± 16.19); *t(*20) = 2.40, *p* = 0.0262 (Figure 3a). However, there were no differences in the frequency of entries in the interaction zone for neither non-stress (IZE: 10.00 ± 0.83) nor stress conditions (IZE: 12.10 ± 1.18); *t*(20) = 1.50 (Figure 3d). Nevertheless, this result indicates that exposure to intermittent social stress induces a deficit in social behavior.

**Figure 3.**
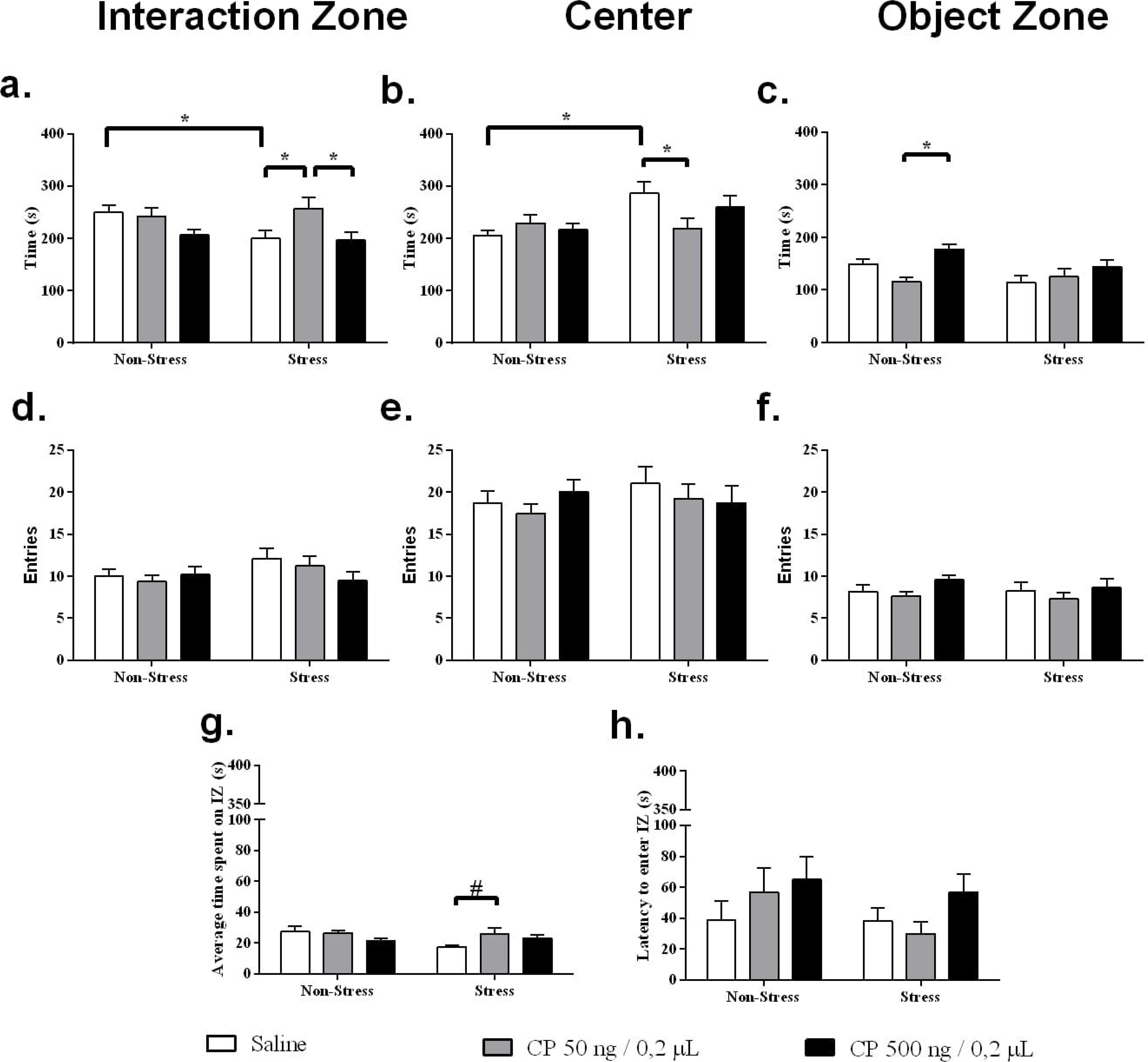
Effects of CP376395, a CRF receptor 1 selective antagonist, administered into the bed nucleus of the stria terminalis of rats exposed to intermittent social defeat stress or non-stressed controls. a. Average time spent in interaction zone; b. Average time spent in the center of the apparatus; c. Average time spent in object zone; d. Average number of entries in interaction zone; e. Average number of entries in the center of the apparatus; f. Average number of entries in object zone; g. Average time spent in interaction zone calculated by the ratio of time spent/frequency of entries in interaction zone; h. Latency to first enter interaction zone. Values were expressed as *M* ± *SEM*; *n* = 11 per group; * *p* < 0.05; # *p* = 0.0508.

To test the effect of CP microinjection combined with the effect of stress exposure, a two-way repeated measures ANOVA was performed. There was a significant interaction between conditions and treatments for time spent in the center of the social interaction apparatus (CeT: *F*(2, 40) = 4.10; *p* = 0.0238). *Post hoc* analysis indicated that this difference was the result of a decrease in time spent in the central chamber for rats from the non-stress condition (CeT: 204.69 ± 9.79) when microinjected with saline compared to rats from the stress condition (CeT: 285.75 ± 22.62); *t(*60) = 3.20, *p* = 0.0061 (Figure 3b). This result was consistent with the initial pairwise comparisons between conditions after saline infusions. No other significant differences for the interaction between conditions and treatments of behaviors displayed in the center of the apparatus were found (IZE: *F*(2, 40) = 1.20; IZT: *F*(2, 40) = 2.50; CeE: *F*(2, 40) = 0.69; CeT: *F*(2, 40) = 0.18; OZT: *F*(2, 40) = 2.20). However, two-way repeated measures ANOVA revealed a significant effect between conditions for time spent in the center (CeT: *F*(1, 20) = 5.00; *p* = 0.0155). *Post hoc* analysis indicated that this difference was related to the decrease of time spent in the central chamber for rats from the stress condition when microinjected with CP 50 ng/0.20 μL (CeT: 218.18 ± 20.27) compared with animals treated with saline (CeT: 285.75 ± 22.62) (Figure 3b). No other significant differences between conditions were found (IZE: *F*(1, 20) = 1.90; IZT: *F*(1, 20) = 1.10; CeE: *F*(1, 20) = 0.47; OZE: *F*(1, 20) = 0.38; OZT: *F*(1, 20) = 4.30). For the effects of drug treatment, two-way repeated measures ANOVA revealed a significant difference between treatments for time spent in the interaction zone (IZT: *F*(2, 40) = 5.30; *p =* 0.0091). *Post hoc* analysis indicated that this difference was related to the increase in time spent in the interaction zone for rats from the stress condition when microinjected with CP 50 ng/0.20 μL (IZT: 256.89 ± 21.80) compared to saline treatment (IZT: 199.74 ± 16.19); *t(*40) = 2.70, *p* = 0.0278, and treatment with CP 50 ng/0.20 μL (IZT: 256.89 ± 21.80) compared to treatment with 500 ng/0.20 μL (IZT: 196.11 ± 15.21); *t(*40) = 2.90, *p* = 0.0177 (Figure 3a).

Surprisingly, two-way repeated measures ANOVA revealed a significant effect between treatments for time spent in the object zone (OZT: *F*(2, 40) = 6.10; *p* = 0.0047). *Post hoc* analysis indicated that this difference was related to the increase of time spent in the object zone for rats from the non-stress condition when microinjected with CP 500 ng/0.20 μL (OZT: 177.25 ± 9.67) compared to treatment with CP 50 ng/0.20 μL (OZT: 115.86 ± 8.81); *t(*40) = 3.70, *p =* 0.0021 (Figure 3c). No other significant differences between treatments were found (IZE: *F*(2, 40) = 1.73; CeE: *F*(2, 40) = 0.45; CeT: *F*(2, 40) = 0.92; OZE: *F*(2, 40) = 2.00).

Next, we examined whether there were differences in the latency to first enter the interaction zone and for how long animals interacted, in an attempt to understand the pro-social effect induced by intra-BNST CP microinjection. Two-way repeated measures ANOVA revealed a trend towards statistical significance on the interaction between condition and treatment for average time spent in the interaction zone (*F*(2, 40) = 3.20, *p* = 0.0508). *Post hoc* analysis indicated that this strong trend was related to the greater average of time spent in the interaction zone for rats from the stress condition when microinjected with CP 50 ng/0.20 μL (25.93 ± 4.05) compared to treatment with saline (17.48 ± 1.15); *t(*40) = 2.5, *p* = 0.0328 (Figure 3g). There were no significant effects on latency to enter the interaction zone between conditions and treatments as revealed by two-way repeated measures ANOVA (*F*(2, 40) = 0.58). The data did not show significant differences in latency between conditions (*F*(1, 20) = 1.50), and did not show significant differences across treatments (*F*(2, 40) = 1.70) (Figure 3h).

## 4. Discussion

In this study, the microinjection of a CRFR1 antagonist into the BNST was able to restore this stress-induced social impairment. The intra-BNST microinjection of CP resulted in an increase of time spent in the interaction zone, as well as longer periods per visit to the compartment containing an unfamiliar conspecific. This implicates the CRF system in the modulation of social behaviors. We did not find differences in the preference for sweet solution across sessions and between non-stressed and stressed rats. This effect was interpreted as a lack of motivational deficits that could otherwise explain the decreased social interaction induced by exposure to social defeat. The CRF system is being profoundly implicated as the link between stress exposure and development of neuropsychiatric conditions [19,20].

We did not find differences between the stress and non-stress conditions for behaviors displayed in the EPM test. This result calls into question the association of intermittent exposure to social defeat with the development of long-lasting anxiety-related symptoms. The EPM test was performed 12 days after the last exposure to social defeat. It is quite possible that the effects of this exposure on the anxiety-associated behaviors analyzed in EPM could be seen at an earlier period. The time interval between stressor exposure and behavioral assessment used in this study gave us the possibility to evaluate longer-lasting stress effects. Another consideration that should be mentioned is that anesthetizing and conducting surgery on the rats could also disrupt the persistence of general anxiety-like behavior.

It is worth remembering that social impairment goes beyond anxiety and is a characteristic symptom of several neuropsychiatric conditions such as autism spectrum disorders, schizophrenia, and stress-adjustment disorders [24–28]. Such disorders present either a possible etiology associated with stress or present aggravated symptoms in situations of exposure to adverse situations [29–31]. Since the characterization of the CRF receptors and its binding protein, CRF-related components are a new class of potential targets for the development of new antidepressants and anxiolytic drugs [21–23]. Strong evidence points to the possibility that CRFR1 is the main receptor that mediates the effects of CRF [7,32]. This assumption is based on binding assays that demonstrate a higher affinity of CRFR1 to CRF than to CRFR2 [33]. Moreover, CRFR2 appears to be the endogenous ligand of urocortins, presenting higher affinity to these CRF-like peptides than to CRF neuropeptide itself [33]. These receptor assays are consistent with several studies conducted with specific CRFR1 antagonists. These studies demonstrate increased anxiety-related behaviors of transgenic mice that overexpress CRF [34]. The first developed CRFR1 selective antagonists demonstrated the ability to inhibit the anxiogenic effects of CRF: NBI27914, CRA1000, CRA1001, and CP154526 [35,36]. Also, anxiolytic effects have been reported with a new class of CRFR1 antagonists, R121919, antalarmin, DMP696 and DMP904 [37–39]. The exact role of both CRFRs is not completely understood.

It has been suggested that the central amygdala and BNST participate in functionally distinct stress-response systems. Rapid excitatory transmission in the central amygdala would be responsible for rapid-onset short-duration threat responses, while activity in the BNST would engender slowly sustained responses to threatening stimuli [40,41]. The processing of a given stressor in respect to its emotional component requires the integration of sensory inputs with prior experiences to determine the salience/valence of the perceived threat. The medial prefrontal cortex, hippocampus, amygdala, and BNST mediate emotional processing and are known to project and stimulate the initial response of the hypothalamic-pituitary-adrenal axis [42]. In this sense, BNST would be responsible to modulate stress-initiating and stress-attenuating responses directed to diffuse and long-onset threatening stimuli, thus the fine balance between these responses seems to be crucial for maintaining a normal physiological and behavioral response to stress [1]. Possibly, the exposure to stress, injury, and drugs can shift the balance to a pathological state. CRF might be an important candidate in modulating these stress-induced neuroadaptations. This might be the case since these BNST-dependent responses directed to threatening stimuli seem to be especially sensitive to CRF receptor blockade and BNST lesions [3]. Therefore, it is tempting to think that, in our study, the increase in the BNST CRF content due to stress exposure might trigger social avoidant behaviors, while the antagonism of CRFR1 in this region might be responsible for preventing this outcome. Although we do not measure the BNST CRF content in this study, there is evidence of CRF mRNA upregulation in the BNST of socially stressed mice [43].

Our results revealed an increase of time spent in the object zone for rats from the non-stress condition when microinjected with the higher dose of CP (500 ng/0.20 μL). This was an unexpected result for two reasons. First, social avoidance is the opposite outcome anticipated for this treatment. Second, CRFR1 antagonists are typically reported as having no activity under baseline conditions in most animal models of stress [44].

Much remains to be unraveled about CRF role in the neurobiology of mood disorders and stress-related psychiatric conditions. The drawbacks presented during clinical testing of CRFR1 antagonists [6,45] increase the understanding of CRF modulation as a narrower pharmacological tool than projected. In this study, exposure to intermittent social stress caused a deficit in social behaviors, evidenced by a lower time spent in the interaction zone of the three-chamber social test. This was observed in the stress group in comparison to the non-stress group when microinjected intra-BNST with saline solution. The microinjection of CP in the BNST was able to restore this stress-induced effect. The results demonstrated that the animals from the stress group spent more time in the interaction zone when microinjected with CP. This was achieved with the microinjection of CP 50 ng/0.20 μL compared to saline. CP produced a dose-dependent effect, resembling an inverted U-shaped dose-response curve, as the microinjection of a higher dose of CP (500 ng/0.20 μL) did not alter the social behavior. This type of dose-response seems to be characteristic among neuropeptide systems [40,46]. These stress-ameliorating effects of CP microinjection were not specific for animals with a previous history of stress. Our results showed that animals from the non-stress group spent more time in the object zone. Although an unexpected result, this could be interpreted as a CP-induced social avoidance in animals from the control group. In conclusion, intra-BSNT CP microinjection restored social behavior of stressed animals to control levels. Indicating that when they were faced with an ambiguous situation, under effect of CP, stressed animals had their preference to spend more time in an environment of social interaction restored.

## Abbreviations

BNST: bed nucleus of the stria terminalis
CeE: Center - Entries
CeT: Center - Time
CP: CP376395
CRF: Corticotropin-Releasing Factor
CRFBP: CRF Binding Protein
CRFR: CRF receptor
EPM: Elevated-plus maze
IZE: Interaction Zone - Entries
IZT: Interaction Zone – Time
OZE: Object Zone - Entries
OZT: Object Zone - Time
PND: Postnatal days

## Acknowledgments

This study was supported by Coordenação de Aperfeiçoamento de Pessoal de Nível Superior (CAPES-Brazil) and Fundo de Incentivo à Pesquisa e Eventos (FIPE-HCPA/UFRGS). L. Albrechet-Souza was supported by CAPES–Brazil (Program CSF-PAJT 88887.096822/2015-00). R. M. M. de Almeida was supported by Conselho Nacional de Desenvolvimento Científico e Tecnológico (CNPq 475176/2012-0). The authors would like to thank Gabriela Jung, Giovana Brum, and Mayra Pachado for assistance in data collection and Fernanda Valiati, Tuane Garcez, Daniela Campagnol, Ana Maria Goldberg, and Marta Cioato for providing helpful technical assistance during the development of this study.

